# *Mycobacterium tuberculosis* genetic features associated with pulmonary tuberculosis severity

**DOI:** 10.1101/2022.05.25.493361

**Authors:** Charlotte Genestet, Guislaine Refrégier, Elisabeth Hodille, Rima Zein-Eddine, Adrien Le Meur, Fiona Hak, Alexia Barbry, Emilie Westeel, Jean-Luc Berland, Astrid Engelmann, Isabelle Verdier, Gérard Lina, Florence Ader, Stéphane Dray, Laurent Jacob, François Massol, Samuel Venner, Oana Dumitrescu, the Lyon TB study group

## Abstract

*Mycobacterium tuberculosis* (Mtb) infections result in a wide spectrum of clinical presentations but without proven Mtb genetic determinants. Herein, 234 pulmonary tuberculosis (TB) patients were stratified according to TB disease severity and Mtb genetic features were explored using whole genome sequencing, including heterologous single nucleotide polymorphism (SNP) calling to explore micro-diversity. Clinical isolates from patients with mild TB carried mutations in genes associated with host-pathogen interaction, while those from patients with moderate/severe TB carried mutations associated with regulatory mechanisms. Genome-wide association study identified a SNP in the promoter of the gene coding for the virulence regulator EspR associated with moderate/severe disease. Structural equation modelling and model comparisons indicated that TB severity was associated with the detection of Mtb micro-diversity within clinical isolates and to the *espR* SNP. Taken together, these results provide a new insight to better understand TB pathophysiology and could provide new prognosis tool for pulmonary TB severity.

## INTRODUCTION

Tuberculosis (TB) caused by *Mycobacterium tuberculosis* (Mtb) complex remains one of the most prevalent and deadly infectious diseases; there were 10 million new cases worldwide in 2020 which led to 1.5 million deaths ^1^. Mtb infections result in a wide spectrum of clinical outcomes, from latent asymptomatic infection to pulmonary or extra-pulmonary manifestations of disease, with an array of symptoms. Such diversity has been historically attributed to host and environmental factors, while the Mtb complex was previously considered genetically monomorphic ^2^. Many Mtb virulence factors are well described but, to date, there are no proven genetic determinants associated with virulence, disease progression, or severity of TB ^3^. However, some Mtb lineages and sublineages were associated with more severe TB in animal models and also in human population studies, suggesting that Mtb genetic factors can affect TB clinical presentation and severity ^4–6^.

Recent Mtb genomic studies have explored the link between specific Mtb polymorphisms and TB clinical presentation ^7,8^. For instance, several compensatory mutations occurring in drug-resistant Mtb clinical isolates were associated with more extensive lung damage ^7^, and an association was found between TB severity and mutations affecting the expression of some components of the ESX-1 secretion system, a key player in Mtb virulence ^8^. In addition, next generation sequencing (NGS)-based studies have revealed micro-diversity in clinical isolates (within hosts, minor variants coexist rather than a clonal colony), and rapid within-host microevolution of Mtb has been suggested by several studies (presence of minor variants within Mtb clinical isolates longitudinally collected upon TB treatment) ^9–14^. Some of these variants harbour drug-resistance mutations, whilst other carry single nucleotide polymorphisms (SNP) in loci involved in modulation of innate immunity and in Mtb cell envelop lipids ^9–14^. In other bacterial species responsible for chronic infections, micro-diversity has been suggested to impact the outcome and severity of illness, being involved in pathogen adaptation to immune response and treatment pressure ^15–18^. Accordingly, we hypothesised that genetic features of Mtb clinical isolates, such as specific polymorphisms or micro-diversity, may be linked to TB severity.

## RESULTS

### TB cohort characterisation

Among the 234 pulmonary TB patients included in this study, 123 had mild disease and 111 moderate/severe disease according to their Bandim TBscore. The median [interquartile range (IQR)] age of the study population was 35 [25-58] years, and a majority were male (66.2%). Most of the patients originated from Europe or Africa, which is consistent with the local epidemiology ^19,20^. No difference was found in terms of comorbidities according to severity group (**Table 1**).

**Table 1:** Patient characteristics.

|  | **Total population**<br>**n=234** | **Mild-grade**<br>**disease**<br>**n=123** | **Moderate/severe-**<br>**grade disease**<br>**n=111** | **p-value** |
| --- | --- | --- | --- | --- |
| **TB severity score** |  |  |  |  |
| Bandim TBscore | 4 [3-6] | 3 [2-4] | 6 [5-7] | **<0.0001** |
| **Demography** |  |  |  |  |
| Age (years) | 35 [25-58] | 38 [26-60] | 32 [22-54] | **0.0335** |
| Sex (male) | 155 (66.2%) | 85 (69.1%) | 70 (63.1%) | 0.3366 |
| Geographical origin, |  |  |  | 0.1044 |
| Europe | 91 (38.9%) | 56 (45.5%) | 35 (31.5%) |  |
| Africa | 116 (49.6%) | 53 (43.1%) | 63 (56.8%) |  |
| Asia | 23 (9.8%) | 11 (8.9%) | 12 (10.8%) |  |
| America | 4 (1.7%) | 3 (2.4%) | 1 (0.9%) |  |
| **Comorbidities** |  |  |  |  |
| EPTB | 68 (29.1%) | 32 (26.0%) | 36 (32.4%) | 0.3141 |
| HIV | 15 (6.4%) | 7 (5.9%) | 8 (7.2%) | 0.7919 |
| Hepatitis | 24 (10.3%) | 9 (7.4%) | 15 (13.5%) | 0.1378 |
| Diabetes | 26 (11.1%) | 18 (15.0%) | 8 (7.3%) | 0.0941 |
| Immunosuppressive therapy | 21 (9.0%) | 11 (9.2%) | 10 (9.0%) | 1.0000 |
| History of TB | 18 (7.7%) | 9 (7.6%) | 9 (8.2%) | 1.0000 |
| Outcomes |  |  |  | **0.0021** |
| Cured | 218 (93.2%) | 114 (92.7%) | 104 (93.7%) |  |
| Fatal outcome | 8 (3.4%) | 1 (0.8%) | 7 (6.3%) |  |
| Unknown | 8 (3.4%) | 8 (6.5%) | 0 (0%) |  |
| **Nutritional status** |  |  |  |  |
| BMI (kg/m<sup>2</sup>) | 20.2 [17.7-22.4] | 21.3 [20.0-23.9] | 18.0 [16.2-20.4] | **<0.0001** |
| Weight loss (%) | 8.0 [4.5-12.3] | 5.9 [2.9-9.0] | 11.4 [6.9-15.4] | **<0.0001** |
| MUST score | 3 [1-5] | 1 [0-3] | 4 [4-6] | **<0.0001** |
| Serum albumin (g/L) | 30.1 [26.0-36.9] | 34.3 [29.2-40.4] | 28.8 [23.7-31.8] | **<0.0001** |
| Total protein (g/L) | 75 [70-81] | 75 [70-80] | 76 [70-81] | 0.7830 |
| Sodium (mmol/L) | 137 [134-139] | 138 [137-139] | 135 [133-138] | **<0.0001** |
| Potassium (mmol/L) | 4.0 [3.8-4.3] | 4.0 [3.8-4.4] | 4.1 [3.8-4.3] | 0.3829 |
| Chloride (mmol/L) | 103 [100-106] | 104 [102-106] | 102 [99-104] | **<0.0001** |
| **Immune status** |  |  |  |  |
| CRP (mg/L) | 47 [13-98] | 21 [6-78] | 72 [23-110] | **<0.0001** |
| CRP to Albumin ratio | 16.6 [4.0-38.7] | 6.6 [1.7-28.9] | 27.4 [9.3-45.1] | **<0.0001** |
| Haemoglobin (g/L) | 120 [104-137] | 130 [116-141] | 111 [97-128] | **<0.0001** |
| White blood cells (G/L) | 6.9 [5.3-9.0] | 6.5 [4.9-8.5] | 7.5 [6.0-9.8] | **0.0071** |
| Neutrophils (G/L) | 4.6 [3.2-6.2] | 4.1 [2.8-5.5] | 5.1 [3.8-6.6] | **0.0006** |
| Eosinophils (G/L) | 0.09 [0.03-0.18] | 0.12 [0.06-0.22] | 0.075 [0.02-0.14] | **0.0003** |
| Basophils (G/L) | 0.03 [0.02-0.05] | 0.03 [0.02-0.05] | 0.03 [0.01-0.05] | 0.3824 |
| Monocytes (G/L) | 0.62 [0.46-0.84] | 0.58 [0.41-0.76] | 0.70 [0.49-1.03] | **0.0061** |
| Lymphocytes (G/L) | 1.41 [0.89-2.01] | 1.54 [0.95-2.14] | 1.23 [0.87-1.63] | **0.0057** |
| Neutrophil to lymphocyte ratio | 3.5 [1.9-5.5] | 2.9 [1.5-4.5] | 4.5 [2.7-6.8] | **<0.0001** |
| Monocyte to Lymphocyte ratio | 0.48 [0.29-0.71] | 0.40 [0.26-0.59] | 0.58 [0.40-0.83] | **<0.0001** |
| Lymphocyte to CRP ratio | 31.2 [11.5-138.3] | 76.9 [18.3-286.4] | 17.9 [9.3-48.1] | **<0.0001** |

Data were expressed as count (%) for dichotomous variables and as median [interquartile range]
for continuous values. The number of missing values was excluded from the denominator. For
dichotomous variables, Fisher's exact or $\chi^2$ test was used as appropriate. For continuous values, non-parametric Mann-Whitney U test or unpaired t-test was used to compare groups as appropriate according to Shapiro-Wilk normality test. *p-value* < 0.05 was considered significant. EPTB: extrapulmonary tuberculosis; unknown outcome: loss to follow-up or follow-up in another care facility; HIV: human immunodeficiency virus; MUST: malnutrition universal screening tool; BMI: body mass index; CRP: C-reactive protein.

As expected, the rate of fatal outcome was more frequent in the moderate/severe-grade group. A poorer nutritional status was also observed in this group, including lower median body mass index (BMI), greater median unintentional weight loss, higher median malnutrition universal screening tool (MUST) score, as well as lower median serum albumin, sodium, and chloride levels. Regarding the immune status, the median level of serum C-reactive protein (CRP), CRP to albumin ratio, as well as white blood cell, neutrophil and monocyte counts, and neutrophil to lymphocyte and monocyte to lymphocyte ratios were higher in the moderate/severe grade group. Conversely, the median haemoglobin level, eosinophil, and lymphocyte counts, as well as the lymphocyte to CRP ratio were lower in this group (**Table 1**).

The proportion of smear-positive patients was higher in the moderate/severe-grade group and accordingly the median time to positivity (TTP) of Mtb cultures was lower (**Table 2**). For both groups, Mtb isolates genetic diversity (**Table 2, Fig. 1** and **Extended Data Fig. 1**) reflected the local epidemiology ^20^. No difference was observed between groups regarding Mtb resistance profile (**Table 2**). Nevertheless, the proportion of Mtb isolates for which micro-diversity was detected (α-diversity >1) was higher in the moderate/severe-grade group, but without difference in the median magnitude of α-diversity (**Table 2**). Of note, no association was observed between Mtb isolate α-diversity magnitude and smear results (**Extended Data Fig. 2A**) or the TTP of Mtb cultures (**Extended Data Fig. 2B**).

**Table 2:** Microbiological characteristics of Mtb isolates.

|  | **Total population**<br>**n=234** | **Mild-grade disease**<br>**n=123** | **Moderate/severe-**<br>**grade disease**<br>**n=111** | ***p-values*** |
| --- | --- | --- | --- | --- |
| Type of sample |  |  |  | 0.0696 |
| Bronchial aspiration | 63 (27%) | 41 (33%) | 22 (20%) |  |
| Biopsy | 10 (4%) | 6 (5%) | 4 (4%) |  |
| Sputum | 137 (59%) | 61 (50%) | 76 (68%) |  |
| BAL | 16 (7%) | 10 (8%) | 6 (5%) |  |
| Stomach tube | 8 (3%) | 5 (4%) | 3 (3%) |  |
| Smear-positive isolates | 111 (47%) | 48 (39%) | 63 (57%) | **0.0087** |
| TTP (days) | 10.5 [6-17] | 12.5 [7-18] | 8 [5-15] | **0.0022** |
| Lineages |  |  |  | 0.2816 |
| L1 | 8 (3%) | 3 (2%) | 5 (4%) |  |
| L2 | 32 (14%) | 18 (15%) | 14 (13%) |  |
| L3 | 13 (6%) | 10 (8%) | 3 (3%) |  |
| L4 | 169 (72%) | 87 (71%) | 82 (74%) |  |
| *M. africanum* | 5 (2%) | 1 (1%) | 4 (4%) |  |
| *M. bovis* | 7 (3%) | 4 (3%) | 3 (3%) |  |
| Resistance profile |  |  |  | 0.2898 |
| Susceptible | 193 (82%) | 97 (79%) | 96 (87%) |  |
| INH monoR | 11 (5%) | 6 (5%) | 5 (5%) |  |
| RIF monoR | 2 (1%) | 1 (1%) | 1 (1%) |  |
| PZA monoR | 1 (0.4%) | 0 (0%) | 1 (1%) |  |
| MDR | 27 (11%) | 19 (15%) | 8 (7%) |  |
| α-diversity >1 | 123 (53%) | 38 (31%) | 85 (77%) | **<0.0001** |
| Magnitude of α-diversity >1 | 1.89 [1.51-2.19] | 1.92 [1.55-2.33] | 1.87 [1.49-2.05] | 0.4495 |

Data were expressed as count (percentage, %) for dichotomous variables and as median [interquartile range] for continuous values. For dichotomous variables, Fisher's exact or $\chi^2$ test was used as appropriate. For continuous values, non-parametric Mann-Whitney U test was used to compare groups. *p-value* < 0.05 was considered significant. BAL: bronchoalveolar lavage;
TTP: time to positivity of Mtb culture. INH: isoniazid, RIF: rifampicin; PZA: pyrazinamide; monoR: mono-resistant; PZA monoR: Mtb isolates resistant to PZA excluding *M. bovis*; MDR: multi-drug resistant.

**Figure 1:**
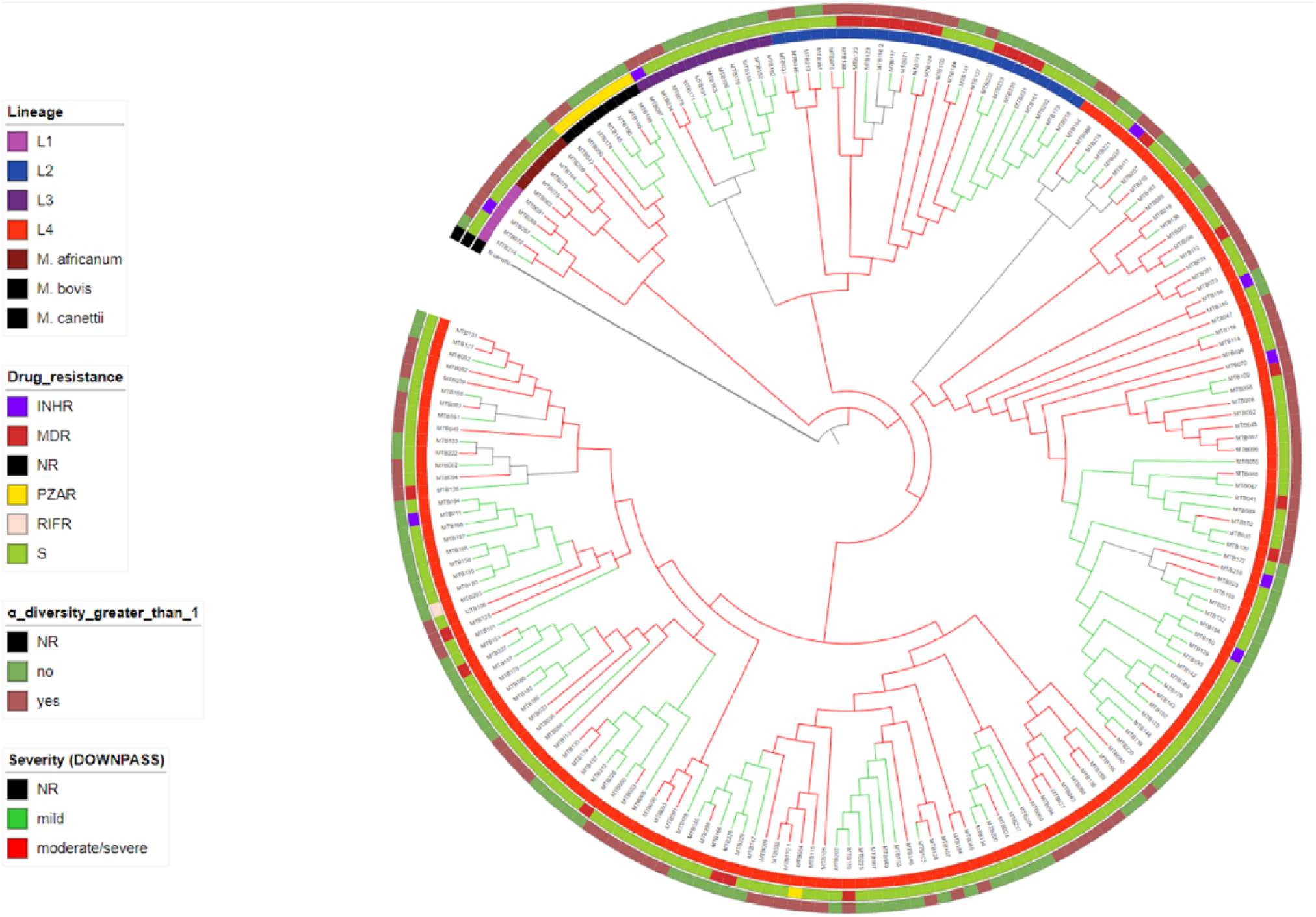
TB severity profile of studied samples and inference of its evolution along the Mtb phylogeny. The phylogeny was reconstructed by Maximum Likelihood that identified all branches with a strong bootstrap support (Extended Data Fig. 1). Tree is displayed with arbitrary branch length to improve visibility. Green branches: mild grade TB severity group; red branches: moderate/severe grade TB severity group. From the inside, rings are coloured by Mtb lineages, drug resistance profile, and occurrence of micro-diversity within clinical isolate (see legend). INHR: isoniazid mono-resistant, RIFR: rifampicin mono-resistant; PZAR: pyrazinamide mono-resistant; MDR: multi-drug resistant; NR: not reported

### Mtb genetic characteristics according to TB severity

We explored the distribution of TB severity profile along the Mtb phylogeny of the strains identified in the present study. Both mild and moderate/severe grade severity profiles were found in several sublineages of each lineage, supporting the inference that this feature evolved recurrently along with Mtb evolution (**Fig. 1**).

To detect Mtb genetic adaptation according to TB severity, we explored polymorphisms in terminal branches of the phylogeny of Mtb samples and within the micro-diversity of Mtb samples through unfixed mutations, both suggestive of ongoing adaptation.

Previous studies have suggested that severe symptoms are associated to a mutational signature typical of oxidative damage (increased changes C > T and G > A) ^21^. Then the precise distribution of Mtb mutations was explored between the mild grade and the moderate/severe grade groups (**Fig. 2**). Differences were observed in the distribution of mutation in the terminal branches, *i*.*e*. fixed mutations of the phylogeny (**Fig. 2A and B**), with a slightly stronger ROS mutational signature in the moderate/severe grade group (*p*<0.0001; **Fig. 2C**), but not within the micro-diversity of Mtb isolates (*p*=0.3132; **Fig. 2D-F**).

**Figure 2:**
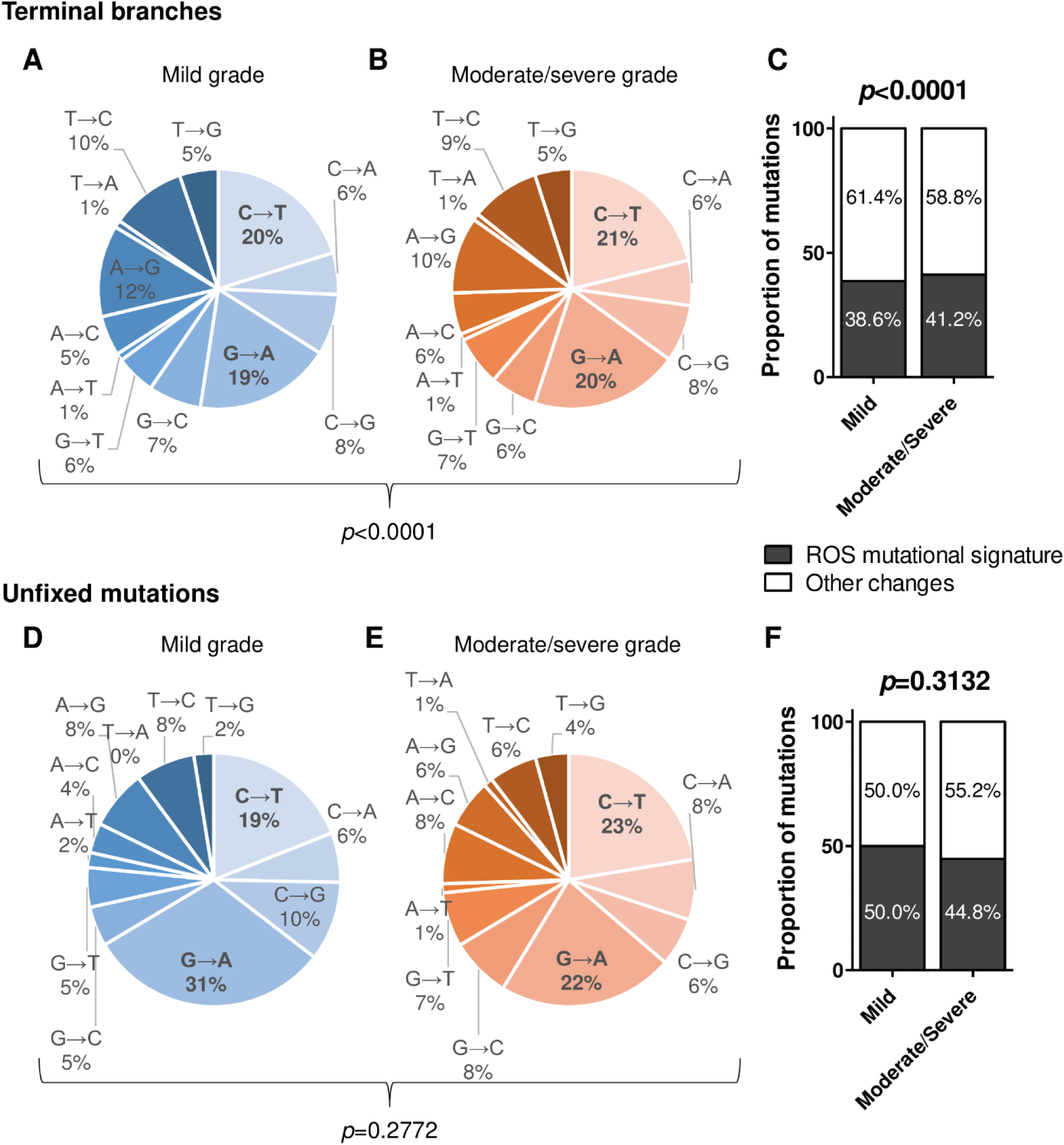
ROS mutational signature according to TB severity. Distribution of mutations in terminal branches of the phylogeny of Mtb samples (A-C; n=21754 mutations explored) and in unfixed mutations within Mtb clinical isolates (D-F; n=437 mutations explored) in the mild-severity grade (A and D) and the moderate/severe-grade groups (B and E). Each type of mutation was explored (A-B and D-E) and a focus was made on ROS mutational signature (C and F). Fisher’s exact or χ2 test was used to compare mild-severity grade and moderate/severe-grade groups, as appropriate.

We then explored the distribution of these mutations across Mtb gene functional categories. No difference was observed for the distribution of non-synonymous (*p*=0.1614) nor synonymous mutations (*p*=0.4815) across gene functional categories in the terminal branches of the phylogeny (**Fig. 3A**). We explored whether some gene functional categories exhibited signs of differential selection pressure in the mild-grade versus moderate/severe-grade group. In the terminal branches of the phylogeny, the gene functional category “virulence, detoxification, adaptation” exhibited both a higher non-synonymous/synonymous mutation ratio (*p=*0.045; **Fig. 3C**) and a higher dN/dS for Mtb strains from the mild grade group (p=6.6×10^−8^; **Extended Data Table 1**). Regarding unfixed variants within Mtb clinical isolates, a difference was observed in the distribution of non-synonymous mutations across gene functional categories (*p*=0.0238) but not in the distribution of synonymous mutation (*p*=0.9019, **Fig. 3B**). The gene functional category “cell wall and cell processes” exhibited both a higher non-synonymous/synonymous mutation ratio and a higher dN/dS in the mild grade group, and the gene functional category “regulatory proteins” did so in the moderate/severe grade group (**Fig. 3D and Extended Data Table 2**).

**Figure 3:**
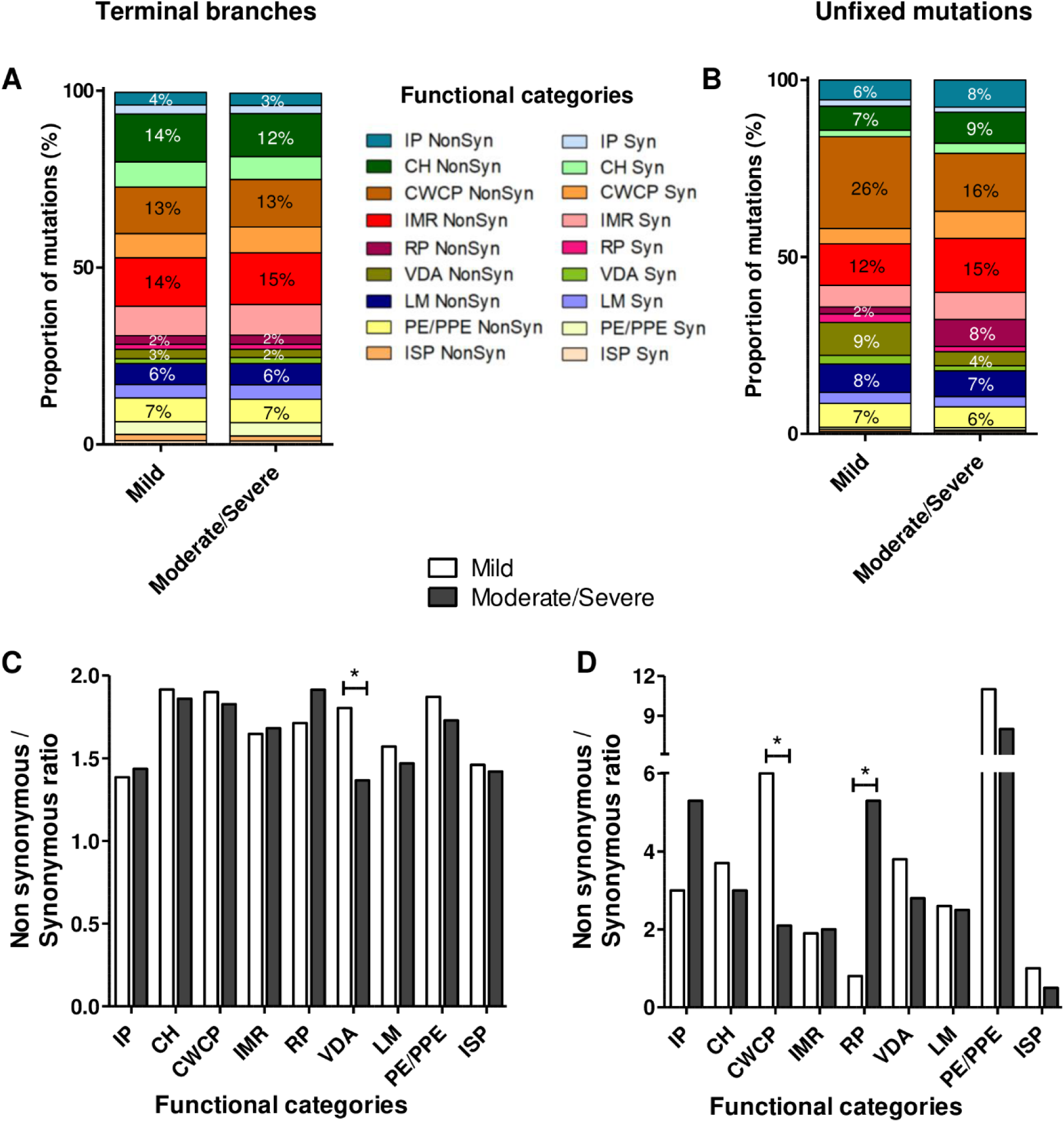
Non-synonymous/synonymous mutations ratio in the gene functional categories according to TB severity. Distribution of non-synonymous and synonymous mutations (A-B) and non-synonymous/synonymous mutation ratio (C-D) across gene functional categories in the terminal branches of the phylogeny of Mtb samples (A-C; n=21729 mutations explored) and in unfixed mutations within Mtb clinical isolates (B-D; n=437 mutations explored) in the mild-severity grade and the moderate/severe-grade groups. Fisher’s exact or χ2 test was used to compare mild-severity grade and moderate/severe-grade groups, as appropriate. IP: Information pathways; CH: Conserved hypotheticals; CWCP: Cell wall and cell processes; IMR: Intermediary metabolism and respiration; RP: Regulatory proteins; VDA: Virulence, detoxification, adaptation; LM: Lipid metabolism; ISP: Insertion sequences and phages; NonSyn: Non-synonymous SNP; Syn: Synonymous SNP. *p<0.05

To characterise the potential Mtb genetic determinants of TB severity, we performed a genome-wide association study (GWAS). GWAS identified a SNP, G4323355C, located in the promoter of the gene *espR*, a gene coding for a regulatory protein of the Mtb ESX-1 secretion system. This SNP was more frequent among Mtb isolates from the moderate/severe-grade group (15/110, 13.5%) than the mild-grade group (4/123, 3.3%; *p*=0.0069).

### Structural equation model (SEM) of TB severity

We performed a SEM analysis to relate TB severity to various explanatory variables. We focused on host variables independent of the stage of TB disease and Mtb genetic features, including those identified above, for the association with TB severity. The TB severity was assessed using the Bandim TBscore, as well as the BMI and proportion (%) of unintentional weight loss. As expected, SEM found that severity markers were correlated with each other. None of the host variables explored had an impact on the TB severity. Among Mtb genetic features, the model showed that only the detection of micro-diversity within Mtb clinical isolates affected TB severity (positive estimated standardized coefficient of 0.52 for α-diversity >1; **Fig. 4A**). The importance of all host and Mtb variables evaluated for TB severity was assessed using model comparisons and indicated that the best model was composed of Mtb α-diversity >1 and the presence of the mutation identified by GWAS (**Fig. 4B and Extended Data Fig. 3**).

**Figure 4:**
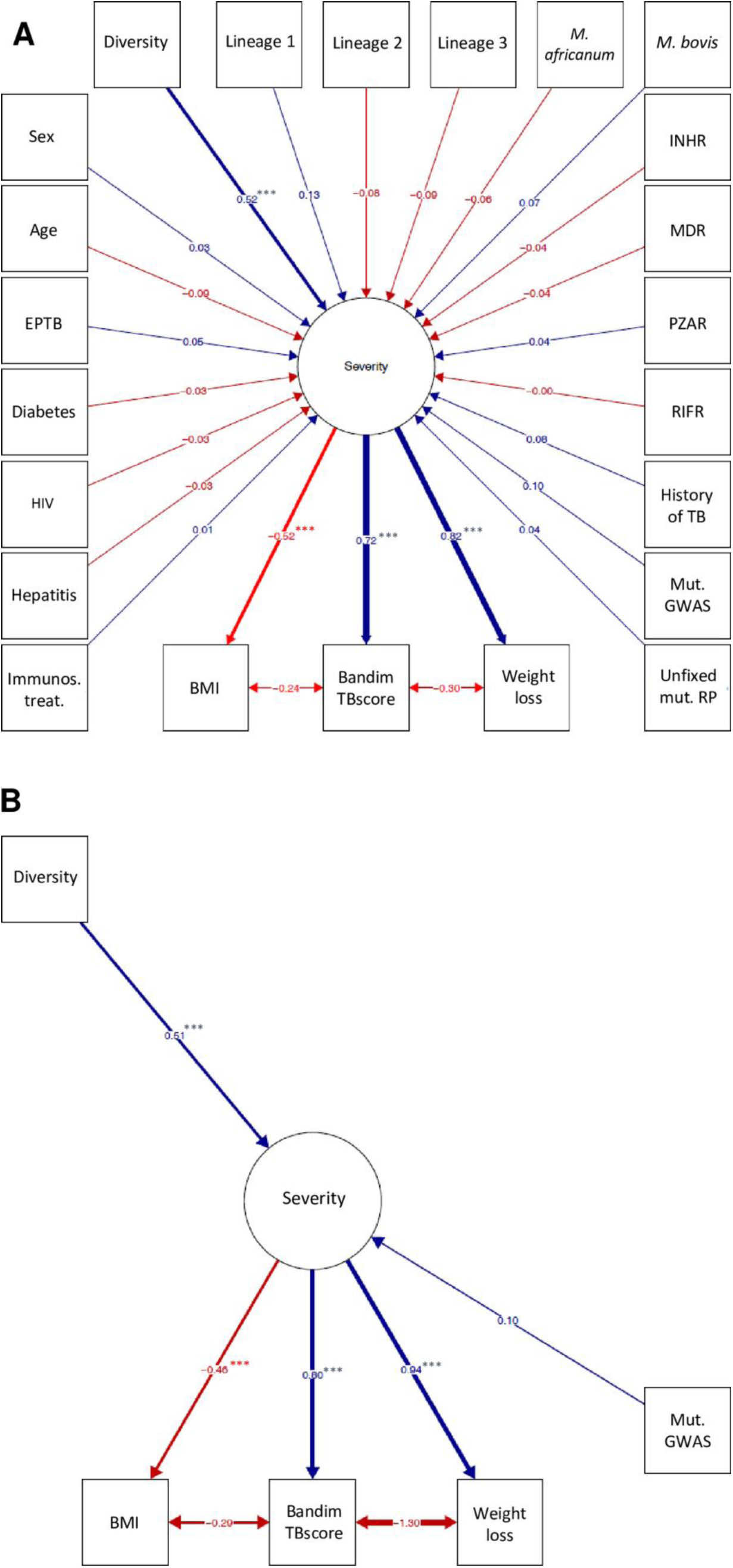
Structural equation modelling (SEM) and the best model examining the effect of host and bacterial factors on pulmonary TB severity. Regarding the variables associated with TB patients, we focused on variables independent of the stage of TB disease, meaning age, sex, ongoing HIV, hepatitis, diabetes, and/or immunosuppressive treatment (Immunos. treat.), history of TB, and extra-pulmonary manifestation (EPTB, both pulmonary and extra-pulmonary). Concerning Mtb genetic features, Mtb lineages (lineage 1, lineage 2, lineage 3, *M. africanum* and *M. bovis*, in contrast with lineage 4), resistance profile (isoniazid mono-resistant [INHR], pyrazinamide mono-resistant [PZAR], rifampicin mono-resistant [RIFR], multidrug resistant [MDR], in contrast with susceptible), detection of Mtb micro-diversity within clinical isolates (Diversity), detection of unfixed mutation in the “regulatory protein” gene functional category (Unfixed mut. RP) and the mutation identified by GWAS (Mut. GWAS) were considered. Unidirectional arrows between variables indicate regression and are associated with standardised regression coefficients. Bidirectional dashed arrows among severity markers indicate correlations. Blue edges indicate positive coefficients or correlations, red edges, negative coefficients or correlations. Residual variance terms are omitted for clarity. *p-value* < 0.05 was considered significant. *p<0.05, ** p <0.01, *** p <0.001. (A) Graphical representation of the maximal model (with all variables). (B) Graphical representation of the best model.

## DISCUSSION

The present study found that Mtb clinical isolates from patients with mild TB carried mutations in genes associated with host-pathogen interaction, while Mtb isolates from patients with moderate/severe TB carried mutations associated with regulatory mechanisms. Moreover, a GWAS-based approach identified a SNP in the promoter of the *espR* gene coding for a regulatory protein, which was found to be associated with TB severity. This SNP is located 1 nucleotide downstream from the transcriptional start site of *espR*, suggesting a potential impact of this mutation on the regulation of this gene; and as the EspR protein regulates 1 of the major Mtb virulence determinants (the ESX-1 system), it would be of interest to further explore the role of this SNP. It is of note that positive selection on other regulatory proteins, such as PhoR, has been reported ^22^, and, taken together, these results point to an overall adaptation to host-pathogen interaction for Mtb strains from patients with moderate/severe disease through regulatory protein involvement.

The finding herein that there was a selection on genes from the “virulence, detoxification, adaptation” functional category, as illustrated by a higher high non-synonymous/synonymous ratio is concordant with that reported previously ^23^. As was the finding that oxidative damage developed upon severe TB disease may be a driver of Mtb diversity ^21^. The integration of the severity status of patients could help to better understand the diverse patterns detected in previous studies exploring the key genetic components involved in sublineage epidemic success ^22,23^. At the same time, in-depth analysis of the impact of Mtb polymorphisms on host-pathogen interaction would help to better understand their involvement in TB pathophysiology.

Furthermore, we applied SEM to identify and evaluate direct and latent interlinkages between, on the one hand, Mtb infection clinical specificities and Mtb isolates’ genetic features and, on the other hand, TB disease severity, to pinpoint the positive and negative influences in this regard. Going beyond the classical linear regression analyses, SEM examines the causal relationships among variables, while controlling simultaneously for measurement error. SEM allowed us to determine the degree of correlation (path coefficients) that capture the importance of a certain path of influence from cause to effect, and it was found that the presence of Mtb micro-diversity within clinical isolates led to greater clinical TB severity. This result needs to be confirmed in an independent prospective validation study.

Previous studies explored the association between Mtb micro-diversity and TB outcome or severity. For instance, the study reported by Nimmo et al. found that Mtb diversity did not affect TB outcome ^11^; this apparent inconsistency with our observations may result from the different read-outs used (outcome versus severity score at time of diagnosis) and different statistical analysis (logistic regression versus SEM). The results presented herein are, however, in accordance with another study that found that greater TB severity was associated with an increase of within-host Mtb micro-diversity, and particularly so in pre-mortem Mtb isolates ^12^. Although we have shown that there is no association between bacterial load and detection of Mtb micro-diversity, it is still unclear whether Mtb micro-diversity is a cause (better Mtb adaptation to treatment, to immune pressure, and/or to various niches) or a consequence (tissue breakdown allowing sampling of Mtb variants usually inaccessible and/or lower immune response reducing selection pressure) of the TB severity.

In addition, no association was found herein between the magnitude of Mtb α-diversity and TB severity. This may be because the analysis was based on the minimum number of variants estimated through whole genome sequencing (WGS) data to calculate Mtb α-diversity, which could lead to underestimate micro-diversity in some Mtb clinical isolate. Nevertheless, detection of unfixed mutations at the level of WGS (meaning mutation frequencies between 10 and 90%) was sufficient to observe a strong association between Mtb micro-diversity detection and TB severity. It is of note that in cancer and microbiological research, calling algorithms for low frequency variants have been developed ^24^ and may be adapted to Mtb WGS data. WGS of Mtb isolates could therefore be envisioned as an all-in-one solution to detect antibiotic resistance ^25^, to infer Mtb transmission chains, to perform epidemiological monitoring ^20,26^, but also as a prognosis tool; the latter would be of value to identify those who would most benefit most from additional management measures, such as therapeutic drug monitoring ^27^.

In conclusion, Mtb micro-diversity within a clinical isolate is related to disease severity. This could be a useful to identify early-on those at high risk of severe TB in order to ensure optimal management.

## METHODS

### Mtb samples, data collection, and ethical considerations

In this single-centre retrospective study, 234 patients diagnosed with microbiologically-proven pulmonary TB from January 2017 to January 2020 at the Lyon University Hospital were included. Moreover, this cohort was enriched with the multidrug resistant (MDR) Mtb cases diagnosed in our centre between June 2010 and December 2016 ^20,26^. For all Mtb clinical isolates, WGS analysis was performed in routine practice as part of the laboratory diagnosis. Demographic (age, sex, continent of birth), clinical (pulmonary, extra-pulmonary TB, symptoms, clinical findings, comorbidities [previous history of TB, active hepatitis, HIV, diabetes, and immunosuppressive treatment at time of TB diagnosis]), microbiological (sputum smear results, time to positivity, antibiotic resistance, lineage, data from WGS), nutritional, and immune data were collected. Only variables for which data was available for ≥80% of patients between 2 weeks before TB diagnosis and 1 week after initiation of anti-TB treatment or nutritional supplementation were considered. Outcomes (cured, fatal outcome, and loss to follow-up) were evaluated 1 year after the end of anti-TB treatment. All data were collected in a database, in accordance with the decision 20-216 of the ethics committee of the Lyon University Hospital and French legislation in place at the time of the study (Reference methodology MR-004 that covers the processing of personal data for purposes of study, evaluation or research that does not involve the individual). Relevant approval regarding access to patient-identifiable information are granted by the French data protection agency (*Commission Nationale de l’Informatique et des Libertés*, CNIL).

### TB-associated severity indices

TB-associated severity indices were evaluated at the time of diagnosis, before initiation of anti-TB treatment or nutritional supplementation.

The modified Bandim TBscore considers 5 symptoms (cough, haemoptysis, dyspnoea, chest pain, night sweats) and 5 clinical findings (anaemia, tachycardia, positive finding at lung auscultation, fever, body mass index [BMI] <18 and <16); 1 point is attributed for each aspect and final score is the sum of these. Patients were stratified into 2 severity classes, mild (Bandim TBscore ≤4) and moderate/severe (≥5) ^28^.

The nutritional status of TB patients was also evaluated using the Malnutrition Universal Screening Tool (MUST) that includes 3 variables (unintentional weight loss score [weight loss <5% = 0, weight loss 5-10% = 1, weight loss >10% = 2], BMI [>20.0 = 0, 18.5-20.0 = 1, <18.5 = 2], and anorexia [if yes = 2]) and the final score is the sum of these ^29^.

### Mtb culture

Mtb clinical isolates were processed as previously described ^25^. Mtb genomic DNA extractions were performed after a single round of culture. Biobanked Mtb isolates were inoculated in mycobacterial growth indicator tube (MGIT) until exponential phase before DNA extraction.

### WGS and Illumina data analysis

Genomic DNA of Mtb-positive cultures was purified from cleared lysate and sequenced on NextSeq or MiSeq system (Illumina, San Diego, USA) at the GENEPII sequencing platform of Lyon University Hospital, as previously described ^26^. Reads were mapped using the BOWTIE2 to the Mtb H37Rv reference genome (Genbank NC000962.2) and variant calling was conducted using SAMtools mpileup, as previously described ^26^. A valid nucleotide variant was called if the position was covered by a depth ≥10 reads and a frequency ≥10%. Regions of genes with repetitive or similar sequences were excluded, i.e. regions of *pe, ppe, pks, pps, esx* gene families The reference genome coverage breadth was ≥93% with a mean depth of coverage of ≥50x. Sequences were submitted to the European Nucleotide Archive (ENA) under accession number PRJEB53047.

### Variant assignment and Mtb α-diversity indices

In a previous study, we showed no significant difference in variant detection and frequencies between sequencing on direct samples and after subculture on media used in routine practice ^30^. Moreover, for the present study, 10 isolates were extracted and sequenced twice to evaluate the variability in mutation frequencies between sequencing experiments. In both sequencing experiments, 52 unfixed mutations were detected at similar frequencies (±10%), ranging from 10 to 90% (**Extended Data Fig. 4A**). Accordingly, to identify the minimum number of variants in each Mtb clinical isolate, a variant was defined as an assembly of mutations at frequencies of ±10% as illustrated in **Extended Data Fig. 4B**. Based on that, the α-diversity index for each isolate was calculated by computing the Rao index of diversity taking into account genetic distance among variants. We computed genetic distance among variants applying Sorensen distance on the presence/absence of 437 mutations and consider this information when computing diversity indices. Following Pavoine et al (2016) ^31^, we rescaled the distances prior to the analysis (dividing by the maximum distance) and use equivalent numbers to allow for comparisons of α-diversities among patients. Note that using equivalent numbers implies that α-diversity is equal to 1 when only one variant is present (no diversity).

### Genome-wide association study (GWAS)

Mtb genomes were assembled using SPAdes-3.14.1 with --careful -t 16 --cov-cutoff auto options. DBGWAS 0.5.4 was then run on the 234 pulmonary clinical isolates for which the Bandim TBscore was available. The contigs obtained from the assembly step were used as input, and the Bandim phenotypes (mild grade [Bandim TBscore ≤4] and moderate/severe grade [≥5]), with default options except -nh=3, -SFF=p100, -nb-cores=6, -nc-db=Resistance_DB_for_DBGWAS.fasta-pt-b=uniprot_sprot_bacteria_for_DBGWAS.fasta.

The two latter options allowed nucleotide and protein level annotation of the results using databases that are available from the DBGWAS repository ^32^.

### Phylogenetic analyses

SNP sequence alignment were purged from any non-phylogenetically informative position using goalign (v0.3.5). A phylogenetic tree was computed by maximum likelihood using the GTR model with RAxML-ng (v1.0.3) and the Stakamakis ascertainment correction. Bootstrap was performed to check for phylogenetic robustness using 100 replicates (**Extended Data Fig. 1)**. Inference of TB severity profile along phylogenetic trees was performed using pastml (v1.9.34). Trees were visualised using iTOL (v6) ^33^.

Polymorphisms were explored in terminal branches of the phylogeny (fixed mutations) and within the micro-diversity of Mtb samples (unfixed mutations). On the one hand, the precise distribution of Mtb mutations was explored to identify mutational signature typical of oxidative damage (increased changes C > T and G > A); on the other hand, differential selection pressure analyses at the level of the gene functional categories were conducted by performing a simple count of non-synonymous and synonymous mutations, and by estimating the selective pressure measured as the dN/dS ratio using the Contrast-FEL method (Fixed-Effect site-Level) in the HyPhy package ^34^. Significant differences between selection pressures acting on the 2 groups at the level of the gene functional categories were tested using re-sampling as described by Coscolla et al. ^35^; 30 re-samplings were sufficient to detect significant differences.

### Statistical analysis

#### Univariate analysis

Data were expressed as count (percentage, %) for dichotomous variables and as median (interquartile range [IQR]) for continuous values. The number of missing values was excluded from the denominator. For dichotomous variables, Fisher’s exact or χ2 test was used as appropriate. For continuous values, the non-parametric Mann-Whitney U test or unpaired t-test was used to compare groups as appropriate and according to the Shapiro-Wilk test of normality. Statistical analyses were performed using GraphPad Prism® for Windows version 5.02 (GraphPad Software, La Jolla, CA, USA). *p-value* < 0.05 was considered significant.

#### Structural equation modelling of severity

To gauge the effect of demographic (age, sex) and clinical (HIV, diabetes, hepatitis, immuno-suppressive treatment, previous history of TB, double location of infection) variables, as well as Mtb genetic features (lineage, antibiotic resistance, occurrence of micro-diversity, occurrence of mutations [SNP identified by GWAS or unfixed mutations in the “regulatory protein” gene functional category]), we performed a latent variable structural equation model (SEM) ^36^ linking all of these variables to a latent severity score, assumed to be expressed through 3 markers: the Bandim TBscore, the BMI, and the unintentional weight loss percentage of patients, which are strongly associated with poor prognosis ^37^. We assumed that the Bandim TBscore, expected to be the best marker of severity, was correlated with the other 2 markers. The model was fitted through maximum likelihood using the R package ‘lavaan’ ^38^. The importance of explanatory variables was assessed using model comparison based on the corrected Akaike Information Criterion (AICc) ^39,40^; from model-specific AICc values we deduced variable weights using the sum of Akaike weights of all models including the focal variable, based on classic methods ^41,42^. When representing the results of SEM, we give standardised coefficient values to allow for comparison between explanatory variables.

## ACKNOWLEDGMENTS

The authors thank Philip Robinson (DRS, Hospices Civils de Lyon, Lyon, France) for help with manuscript preparation, the GENEPII sequencing platform (*Institut des agents infectieux*, Hospices Civils de Lyon, Lyon, France) for the MTBC strains sequencing, and the Institute for Integrative Biology of the Cell (I2BC, Université Paris Saclay, Gif-sur-Yvette, France) for the use of their sharing platform.

## EXTENDED DATA

**Extended Data Table 1:**
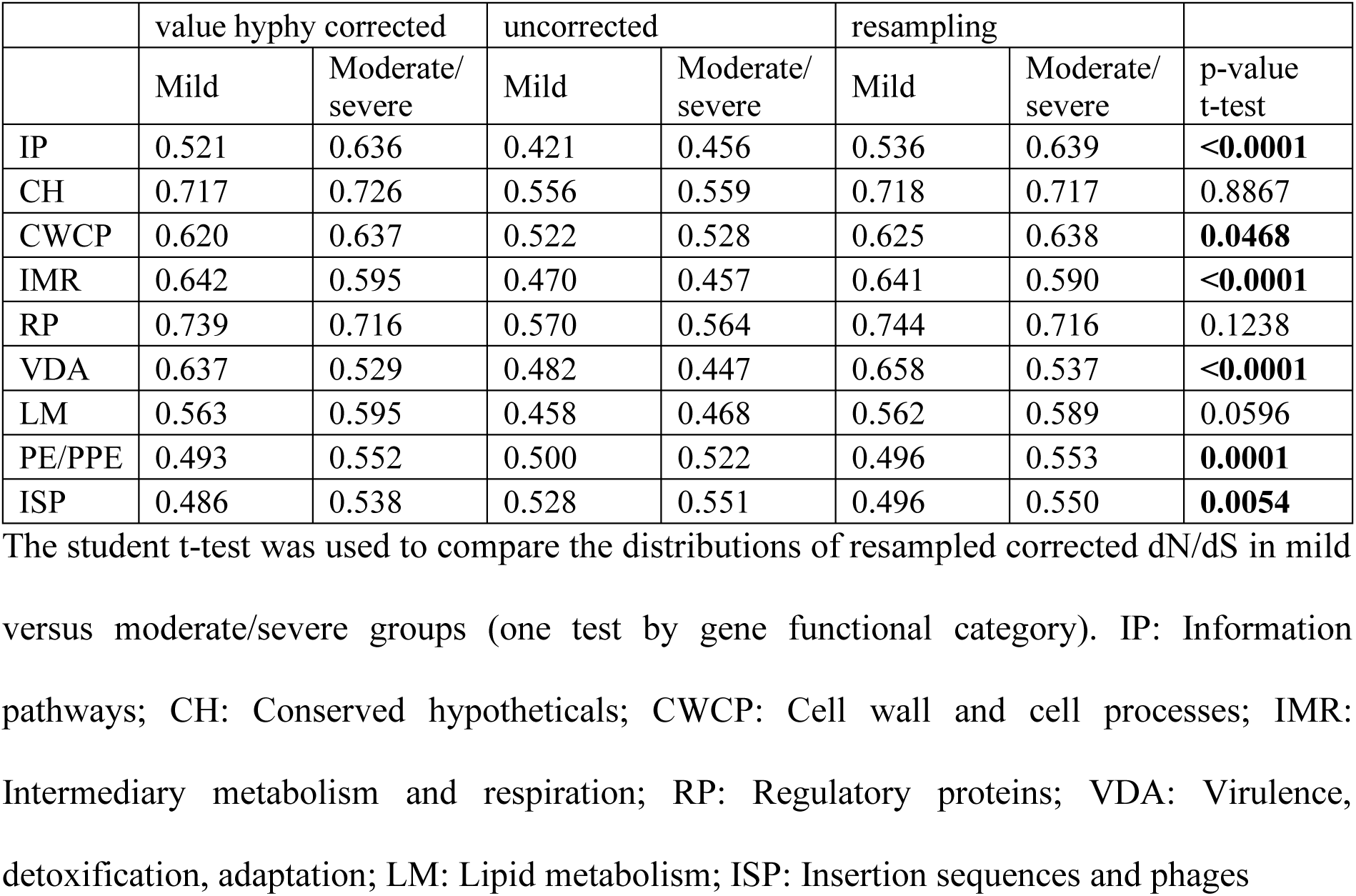
dN/dS ratio across gene functional categories in the terminal branches of the phylogeny.

|  | value hyphy corrected |  | uncorrected |  | resampling |  |  |
| --- | --- | --- | --- | --- | --- | --- | --- |
|  | Mild | Moderate/severe | Mild | Moderate/severe | Mild | Moderate/severe | p-value t-test |
| IP | 0.521 | 0.636 | 0.421 | 0.456 | 0.536 | 0.639 | **<0.0001** |
| CH | 0.717 | 0.726 | 0.556 | 0.559 | 0.718 | 0.717 | 0.8867 |
| CWCP | 0.620 | 0.637 | 0.522 | 0.528 | 0.625 | 0.638 | **0.0468** |
| IMR | 0.642 | 0.595 | 0.470 | 0.457 | 0.641 | 0.590 | **<0.0001** |
| RP | 0.739 | 0.716 | 0.570 | 0.564 | 0.744 | 0.716 | 0.1238 |
| VDA | 0.637 | 0.529 | 0.482 | 0.447 | 0.658 | 0.537 | **<0.0001** |
| LM | 0.563 | 0.595 | 0.458 | 0.468 | 0.562 | 0.589 | 0.0596 |
| PE/PPE | 0.493 | 0.552 | 0.500 | 0.522 | 0.496 | 0.553 | **0.0001** |
| ISP | 0.486 | 0.538 | 0.528 | 0.551 | 0.496 | 0.550 | **0.0054** |

The student t-test was used to compare the distributions of resampled corrected dN/dS in mild versus moderate/severe groups (one test by gene functional category). IP: Information pathways; CH: Conserved hypotheticals; CWCP: Cell wall and cell processes; IMR: Intermediary metabolism and respiration; RP: Regulatory proteins; VDA: Virulence, detoxification, adaptation; LM: Lipid metabolism; ISP: Insertion sequences and phages

**Extended Data Table 2:** dN/dS ratio across gene functional categories within Mtb micro-diversity (unfixed mutations)

|  | value hyphy corrected |  | uncorrected |  | resampling |  |  |
| --- | --- | --- | --- | --- | --- | --- | --- |
|  | Mild | Moderate/severe | Mild | Moderate/severe | Mild | Moderate/severe | p-value t-test |
| IP | 0.373 | 28.180 | 0.421 | 0.845 | 2.237 | 17.406 | **<0.0001** |
| CH | 0.558 | 0.359 | 0.597 | 0.491 | 0.625 | 3.209 | 0.0997 |
| CWCP | 1.395 | 0.942 | 0.753 | 0.685 | 4.866 | 1.192 | **0.0272** |
| IMR | 31.245 | 1.193 | 0.943 | 0.726 | 27.621 | 2.211 | **<0.0001** |
| RP | 0.194 | 0.784 | 0.363 | 0.727 | 1.158 | 8.333 | **0.0080** |
| VDA | 0.365 | 28.971 | 0.450 | 0.901 | 6.970 | 27.371 | **<0.0001** |
| LM | 28.432 | 0.166 | 0.754 | 0.251 | 26.538 | 39.884 | 0.7152 |
| PE/PPE | 27.294 | 26.310 | 0.819 | 0.817 | 23.967 | 21.713 | 0.3910 |
| ISP | 0.000 | 0.833 | 0.000 | 0.914 | 0.344 | 0.847 | **<0.0001** |

The student t-test was used to compare the distributions of resampled corrected dN/dS in mild versus moderate/severe groups (one test by gene functional category). IP: Information pathways; CH: Conserved hypotheticals; CWCP: Cell wall and cell processes; IMR: Intermediary metabolism and respiration; RP: Regulatory proteins; VDA: Virulence, detoxification, adaptation; LM: Lipid metabolism; ISP: Insertion sequences and phages.

**Extended Data Figure 1:**
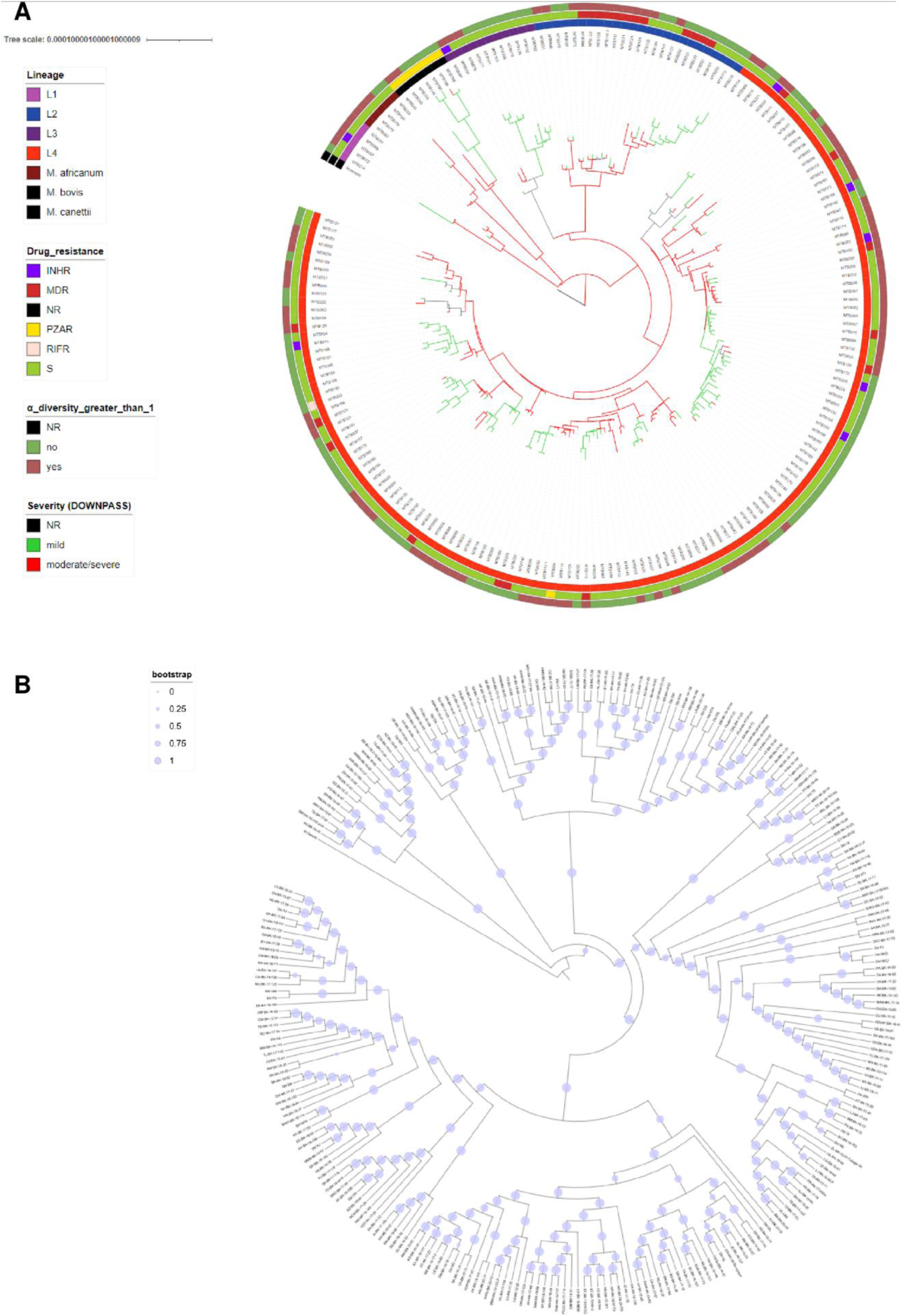
Phylogenetic reconstruction and TB severity profile of studied samples and inference of its evolution along the Mtb phylogeny. **A**. Phylogeny reconstructed by Maximum Likelihood and inference of TB severity profile for each branch. Green branches: mild grade TB severity group; red branches: moderate/severe grade TB severity group. From the inside, rings are coloured by Mtb lineages, drug resistance profile, and occurrence of micro-diversity within clinical isolate (see legend). INHR: isoniazid mono-resistant, RIFR: rifampicin mono-resistant; PZAR: pyrazinamide mono-resistant; MDR: multi-drug resistant; NR: not reported. **B**. Phylogeny showing the bootstrap supports for each branch (100 replicates). Radius of blue discs is proportional to bootstrap support. Almost all branches show a bootstrap support of 100.

**Extended Data Figure 2:**
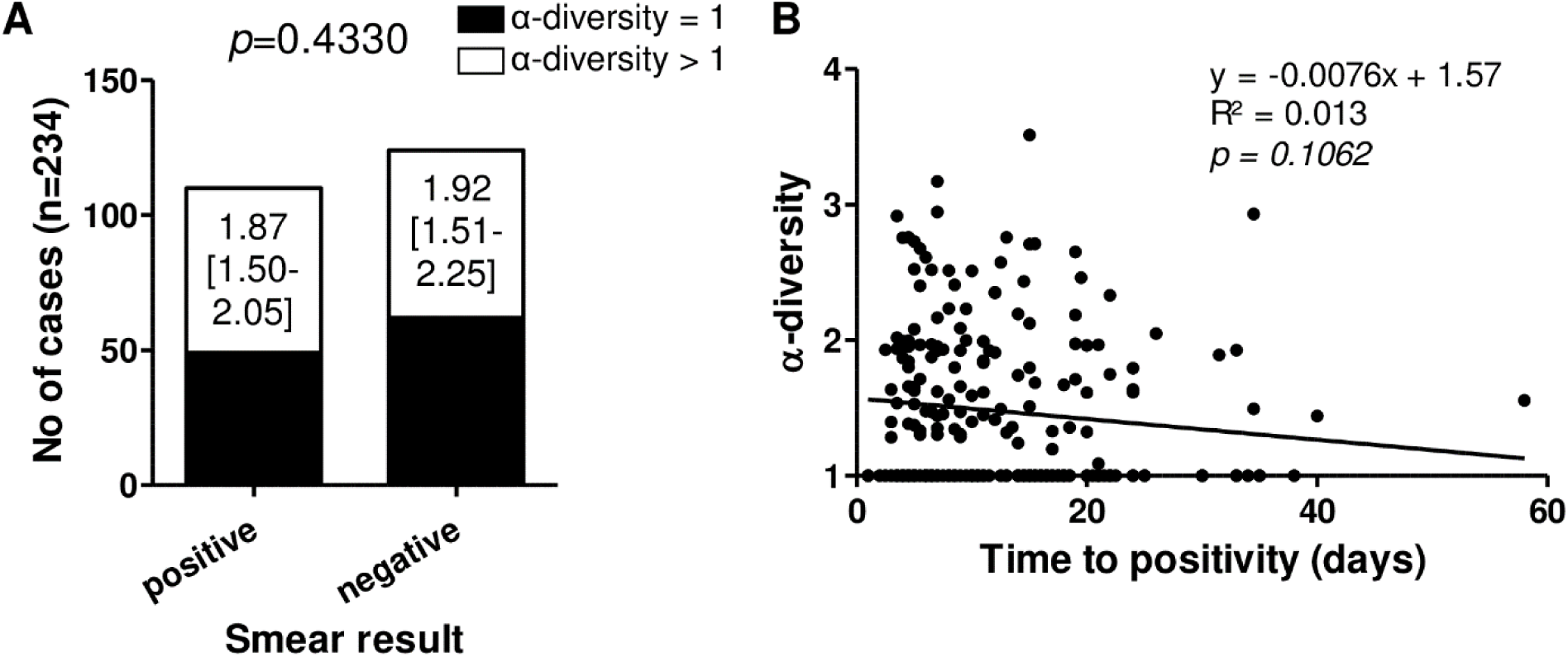
Link between Mtb α-diversity and the bacterial load in Mtb clinical isolates. Association between the detection and the magnitude of Mtb α-diversity and smear results (A) and time to positivity of Mtb clinical isolates (B). A. Black bar: α-diversity=1 no diversity detected by WGS; white bar: α-diversity>1 at least 2 variants detected by WGS. Fisher exact was used to compare groups. x.xx [y.yy-z.zz]: median [IQR] of Mtb α-diversity. Mann-Whitney U test was used to compare magnitude of Mtb α-diversity between groups (no significant differences observed). B. Linear regression between time to positivity of Mtb culture and their respective α-diversities. *p-value* < 0.05 was considered significant.

**Extended Data Figure 3:**
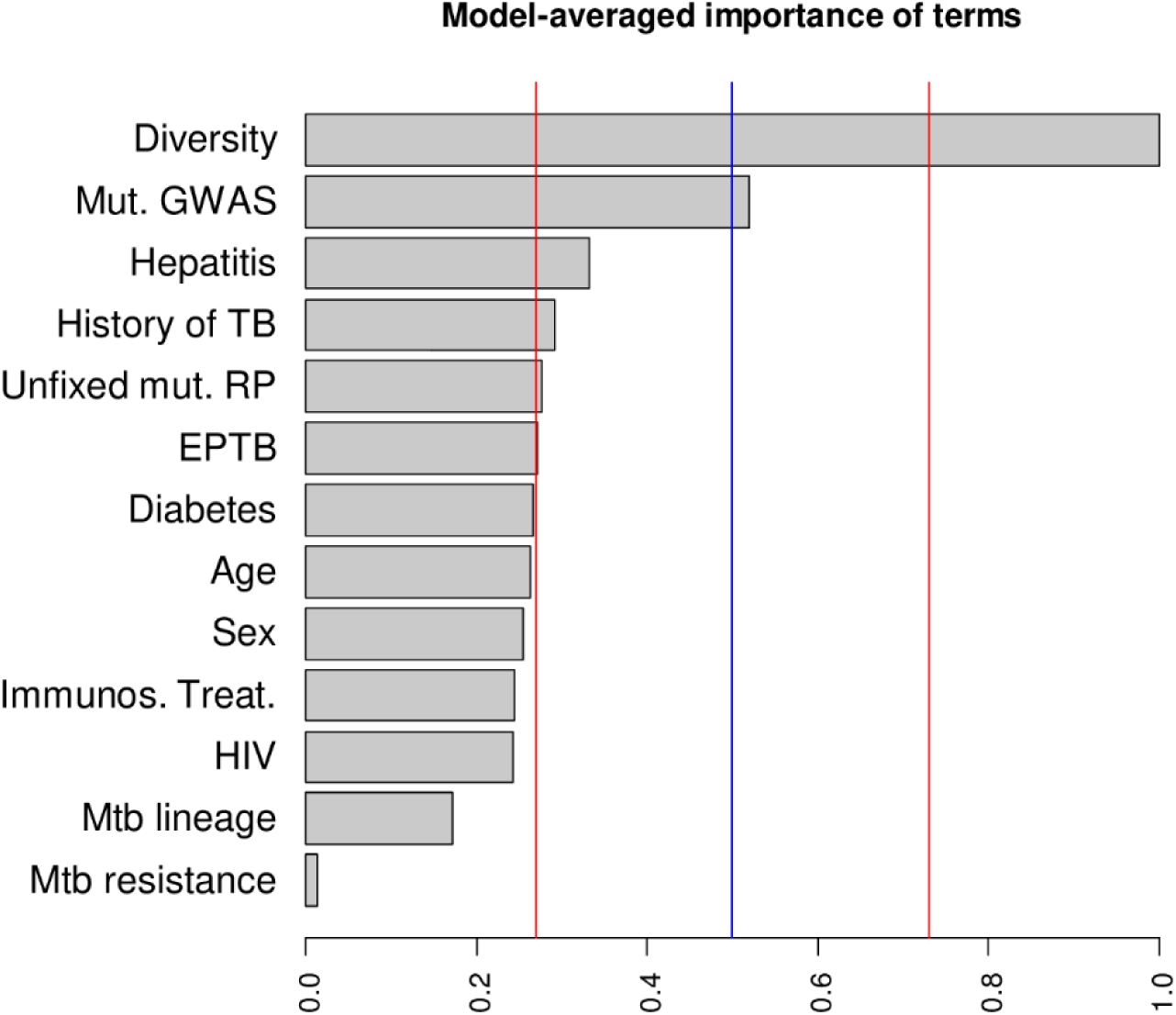
Model-averaged importance of terms for pulmonary TB severity. Model-averaged importance of each term in the model, which is defined as the sum of Akaike weights of models in which a given term appears. Blue line indicates 50% support, i.e. the limit to the plausibility of a term since all variables were included in exactly half of the tested models. Red lines indicate limits between likely and plausible effects (at a sum of weights of 0.73) and between unlikely and implausible ones (at 0.27). The variables considered were: Mtb lineage and resistance profile, detection of Mtb micro-diversity within clinical isolates (Diversity), the mutation identified by GWAS (Mut. GWAS), detection of unfixed mutation in the “regulatory protein” gene functional category (unfixed mut. RP), age, sex, extra-pulmonary manifestation (EPTB, both pulmonary and extra-pulmonary), previous history of TB, and ongoing hepatitis, diabetes, immunosuppressive treatment (Immunos. treat.), and/or HIV.

**Extended Data Figure 4:**
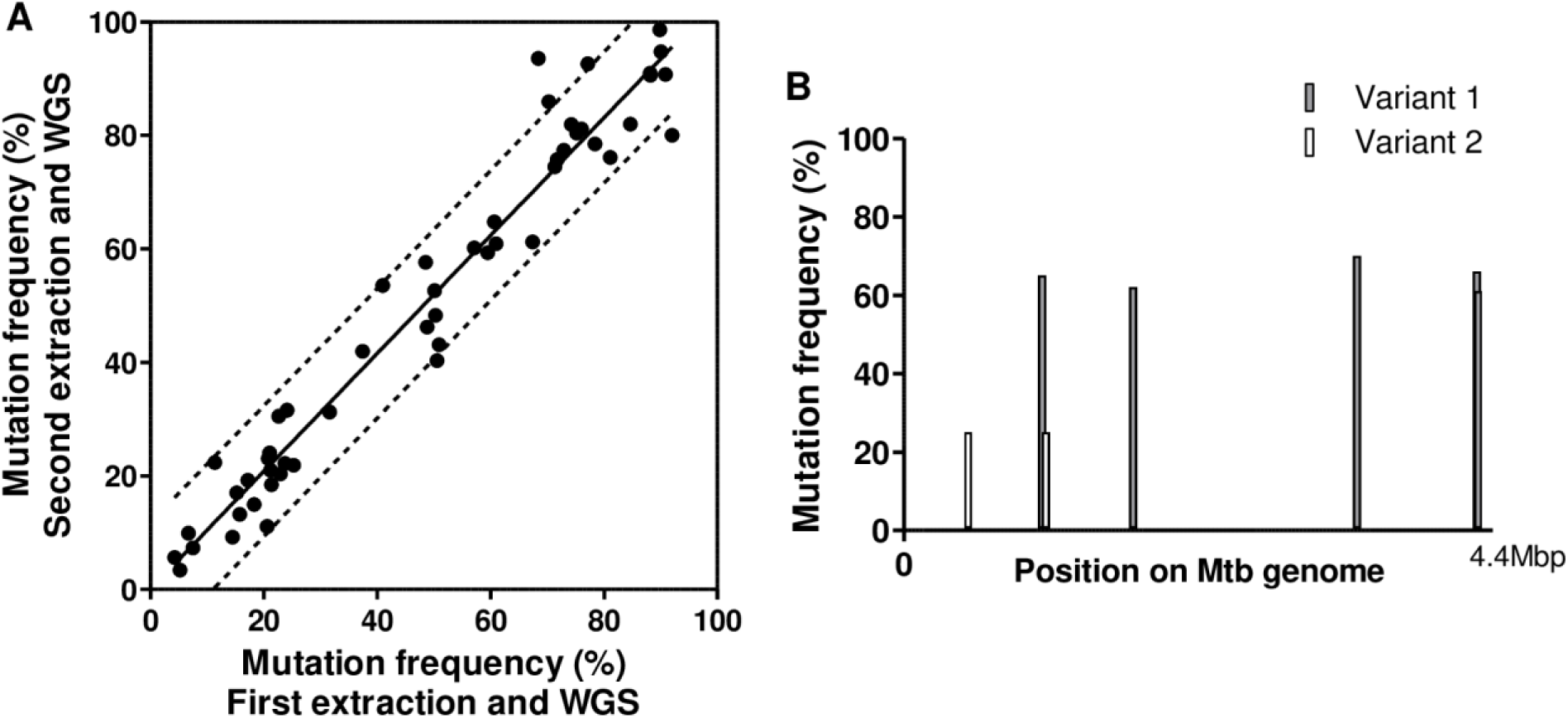
Method for variant assignment in Mtb clinical isolates. A. 10 isolates, containing 52 unfixed mutations, at frequencies ranging from 10% to 90%, were extracted and sequenced twice to evaluate the variability in mutation detection and frequencies between experiments. Continuous line is the linear regression curve. Discontinuous bands are the 90% prediction bands. B. Mutation frequencies across Mtb genome, obtained by WGS, each bar representing a mutation. For variant 1, 5 mutations at frequencies of 65, 62, 70, 66 and 61% were observed, indicating a frequency of 65% for this variant. For variant 2, 2 mutations were observed both at a frequency of 25%, indicating a frequency of 25% for this variant. A third variant, not carrying any of these 7 mutations, is estimated at a frequency of 10%.

